# Timing malaria transmission with mosquito fluctuations

**DOI:** 10.1101/251728

**Authors:** R. Pigeault, Q. Caudron, A. Nicot, A. Rivero, S. Gandon

## Abstract

Temporal variations in the activity of arthropod vectors can dramatically affect the epidemiology and evolution of vector-borne pathogens. Here we explore the “Hawking hypothesis” stating that these pathogens may evolve the ability to time investment in transmission to match the activity of their vectors. First, we use a theoretical model to identify the conditions promoting the evolution of time-varying transmission strategies in pathogens. Second, we experimentally test the “Hawking hypothesis” by monitoring the within-host dynamics of *Plasmodium relictum* throughout the acute and the chronic phases of the bird infection. To explore the periodicity in the host parasite density, we develop a new methodology to correct for non-stationarities in the host parasitaemia. We detect a periodic increase of parasitaemia and mosquito infection in the late afternoon that coincides with an increase in the biting activity of its natural vector. We also detect a positive effect of mosquito bites on *Plasmodium* replication in the birds both in the acute and in the chronic phases of the infection. This study highlights that *Plasmodium* parasites use two different strategies to increase the match between transmission potential and vector availability. We discuss the adaptive nature of these unconditional and plastic transmission strategies with respect to the time-scale and the predictability of the fluctuations in the activity of the vector.

**Impact Summary:** Seasonal and daily fluctuations in the environment affect the abundance and the activity of vectors and may therefore have profound consequences on the transmission of infectious diseases. Here we show that, in accord with evolutionary theory, malaria parasites have evolved two different and complementary strategies to cope with fluctuations in mosquito availability. First, *Plasmodium relictum* adopts an unconditional strategy whereby within-host parasitaemia and mosquito infection increases in the afternoon and in the evening, when its vector, the *Culex pipiens* mosquito, is most active. Second, we find evidence for a plastic strategy allowing the parasitaemia to rapidly increase after exposure to mosquito bites.

## INTRODUCTION

All organisms face periodic changes in their environment. These environmental fluctuations, which can happen at time scales ranging from daily to annual, affect the physiological, immunological and behavioural activities of all species (Smaaland *et al.* 2002; Corder *et al.* 2016; Duboscq *et al.* 2016) including parasites (Martinez-Bakker & Helm 2015; Thaiss *et al.* 2015; Rijo-Ferreira *et al.* 2017a). Both short term (circadian) and long term (seasonal) fluctuations in the environment may trigger dramatic perturbations of the physiology of the hosts that can affect the within host dynamics of the parasite and, ultimately, its epidemiology. One potential explanation for these parasite fluctuations is that they are a by-product of the biological rhythms imposed by the host. There is, for example, abundant evidence of the existence of short-term (circadian) rhythms in the expression of physiological and immune host genes that may potentially impact the development of the parasites within (Edgar *et al.* 2016). Longer-term (seasonal) fluctuations may also trigger dramatic perturbations of the physiology and immunology of the host, which may affect the within-host dynamics of some parasites (see Martinez-Bakker & Helm 2015).

Alternatively, and arguably more interestingly, these periodic fluctuations may be viewed as pathogen adaptations aimed at maximizing transmission by taking advantage of a transient favourable environment (Hawking 1975; Martinez-Bakker & Helm 2015). For instance, in the coccidian parasite *Isospora sp,* the highly synchronized production of transmissible stages in the faeces of infected animals takes place in the late afternoon to minimize mortality through desiccation and UV radiation (Martinaud *et al.* 2009). Crucially, Hawking (Hawking 1970, 1975) argued that similar processes may be acting in vector-borne diseases. He postulated that the timing and the rhythm of many vector-borne pathogens may have evolved to match the daily fluctuations in vector abundance. This so-called “Hawking hypothesis” (Garnham & Powers 1974; Gautret & Motard 1999) has received considerable empirical support from both within and cross-species comparisons of microfilarial parasites, where parasite and mosquito daily rhythms seem to be well matched. For example, the parasite *Wuchereria bancrofti*, which is transmitted by night-biting *Culex sp* mosquitoes, shows a marked nocturnal periodicity where the transmissible microfilaria are sequestered in the lungs during daytime and released into the peripheral blood at night (Hawking 1975). However, in the Pacific islands, where the parasite is transmitted by day-biting *Aedes polinesiensis* mosquitoes, *Wuchereria bancrofti* microfilaria are significantly more abundant during the day (Moulia-Pelat *et al.* 1993).

Many malaria parasites exhibit striking periodic and synchronized cell cycles leading to the simultaneous burst of infected red blood cells at regular points in time. In spite of numerous studies exploring the adaptive nature of malaria periodicity in relation to vector activity (Hawking 1970, 1975; Gautret & Motard 1999) whether these patterns fit the “Hawking hypothesis” remains a controversial issue. Mideo *et al.* (2013) put forward three main arguments against the validity of the Hawking hypothesis in malaria. First, they argued that an accurate timing of gametocyte *production* requires a very finely-tuned synchronization of the whole parasite life cycle. We agree, but contend that gametocyte *maturation* (the process under which gametocytes become infective to mosquitoes, Alano 2007) may be decoupled from the rest of the parasite’s life cycle. Although the process of gametocyte maturation is still not well understood, potential inducers of gametocyte maturation have been described (Sinden 2015), some of which may be under circadian control (Rijo-Ferreira *et al.* 2017b). Second, Mideo *et al.* (2013) also argued that even if the timing of the production of infectious gametocytes is perfectly controlled, the long life expectancy of mature gametocytes would erase any daily rhythm imposed on their production. The gametocytes of most *Plasmodium* species seem, however, to have very short lifespans, surviving for a few hours after their production (see Gautret & Motard 1999, Alano 2007). The one exception is *P. falciparum* whose gametocytes seem indeed to live an inordinate amount of time (6 days, Bousema & Drakeley 2011). Whether these mature gametocytes remain infective throughout their lifespan is, however, not entirely clear. Indeed, the expected positive correlation between gametocyte density and mosquito infection is often not very strong (Bousema & Drakeley 2011). Hawking (1966), for instance, observed that “the cycle of infectivity is not due to the cycle of the *number* of gametocytes in the blood but must be due to variation in their *physiological* state – i.e., their suitability to develop in mosquitoes”. This suggests that malaria infectivity is not driven solely by gametocyte abundance. The last of Mideo *et al.*’s (2013) objections is the lack of evidence for a match between the parasite’s cycles in infectivity and the biting activity of mosquitoes. This is a point we agree with, as the large majority of studies aiming to test the Hawking hypothesis in malaria have indeed focused on the within-host dynamics of the parasite, without testing whether this translates into higher mosquito infection.

Here, we first present a theoretical model that studies evolution of time-varying transmission strategies of *Plasmodium* in a periodically fluctuating environment. This model identifies the conditions under which a periodic investment in transmission is expected to evolve. Then, we carry out an experiment to explore empirically the validity of the “Hawking hypothesis”. For this purpose, we study the periodicity of the avian malaria parasite, *Plasmodium relictum,* in relation with the timing of the activity of its natural vector in the field, the mosquito *Culex pipiens.* In contrast with human malaria, *P. relictum* does not exhibit synchronous development in its vertebrate host (all erythrocytic stages are present in the blood at all times) but several earlier studies report daily fluctuations in within-host parasite abundance (see Gambrell 1937; Hewitt 1940). Yet, the potential link between these fluctuations and the activity of the mosquito vectors remains to be investigated. To explore the validity of the “Hawking hypothesis” we monitored both blood parasitaemia (Pigeault *et al.* 2015) and mosquito activity throughout the day. We use overall parasitaemia as a proxy for transmissible stage (gametocyte) production because in avian malaria the development of gametocytes follows quite closely the development of asexual forms (Hewitt 1940) and our previous work (Pigeault et al 2015) has shown that there’s a very good correlation between sexual (gametocyte) and asexual parasitaemia. As pointed out by our theoretical analysis, the adaptive scenario underlying the “Hawking hypothesis” should yield a positive covariance between bird parasitaemia and mosquito activity.

We worked on both the acute and chronic stages of the infection. From the point of view of the parasite these two stages are fundamentally different in terms of transmission opportunities. While the acute phase is very short lived and results in high rates of mosquito infection, the chronic phase can last several months, and even years, but does not yield high transmission rates (Cornet *et al.* 2014; Pigeault *et al.* 2015). We thus compare these two phases of the infections to establish: (i) the existence of fluctuations of blood parasitaemia throughout the day and (ii) whether these fluctuations translate into higher pathogen transmission to mosquitoes. To explore the periodicity in host parasite density, we developed a new methodology to correct for non-stationarities in the host parasitaemia caused by the large-scale changes in within-host dynamics during the acute phase of the infection. In addition, given that mosquito bites may themselves affect within-host dynamics of the parasite (Lawaly *et al.* 2012; Cornet *et al.* 2014; Reece & Mideo 2014) we compared the within-host dynamics of malaria in birds exposed (or not) to mosquitoes. Mosquito bites may be yet another way for the parasite to respond to the variability of the environment, albeit at a different (shorter) temporal scale. We have previously argued that such a strategy may be an adaptation to a fluctuating seasonal environment where mosquitoes are very abundant during certain seasons and absent during others (Cornet *et al.* 2014; Reece & Mideo 2014). The present paper is an attempt to explore another dimension of malaria adaptation to fluctuations in mosquito availability. In the following we show that *Plasmodium* parasites can use both constitutive and plastic variations in within-host investment in transmission to match short-term and long-term fluctuations in vector availability.

## MATERIAL & METHODS

### Malaria parasites and mosquitoes

*Plasmodium relictum* (lineage SGS1) is the aetiological agent of the most prevalent form of avian malaria which is commonly found infecting passeriform birds in Europe (Pigeault *et al.* 2015). Our parasite lineage (SGS1) was isolated from an infected house sparrow caught in the region of Saintes Maries-de-la-Mer (France) in May 2015 and transferred to naïve canaries (*Serinus canaria,* Passeriforms).

Mosquito experiments were conducted with a laboratory isogenic strain of *Cx. pipiens* mosquitoes. The susceptibility to infection by *P. relictum* and the behavioural activity of our mosquito strain are similar to what is observed in wild *Cx. pipiens* mosquitoes (Vézilier *et al.* 2010, Pigeault *pers. obs.*). Mosquitoes were reared as described by Vézilier *et al.* (2010). We used females 7 ±2 days after emergence that had no prior access to blood and which were starved for 6h before the experiment. Mosquitoes and canaries were maintained under a 12:12-h LD cycle (6h light on, 18h light off).

### Experimental design

Experiments were carried out using (1- year old) domestic canaries (*Serinus canaria*). Prior to the experiments, a small amount of blood (3-5μL) was collected from the medial metatarsal vein of each of the birds and used to verify that they were free from any previous haemosporidian infections. Eight canaries were experimentally inoculated by means of an intraperitoneal injection of *ca.* 80μL of an infected blood pool (day 0, **Fig. 1**, Pigeault *et al.* 2015). The blood pool was constituted of a mixture of blood from 3 infected canaries inoculated with the parasite isolated from the field three weeks before the experiment. The eight infected birds were assigned to two treatments: “exposed” (n=3) or “unexposed” (n=5) to mosquito bites. One “unexposed” bird lost the malaria infection very quickly (10 dpi) and was removed from the analyses. From day 8 to day 70 post-infection parasitaemia of each bird was monitored regularly at noon (12h, **Fig. 1**) except during the experimental sessions when sampling was increased to 4 times per day (see below for details). All blood samples were carried out by collecting 5-10μL of blood from the medial metatarsal vein. A drop of this blood sample was smeared onto a slide for the visual quantification of the parasitaemia (Valkiunas 2004), and the rest was frozen for the molecular quantification of the parasitaemia (see below). In *Plasmodium relictum* infections parasitaemia and gametocytaemia are strongly positively correlated (see Figure 2 in Pigeault *et al.* 2015). For practical reasons, parasitaemia, which is more rapidly quantified, was therefore used as a proxy of parasite investment in the production of transmissible stage.

**Figure 1:**
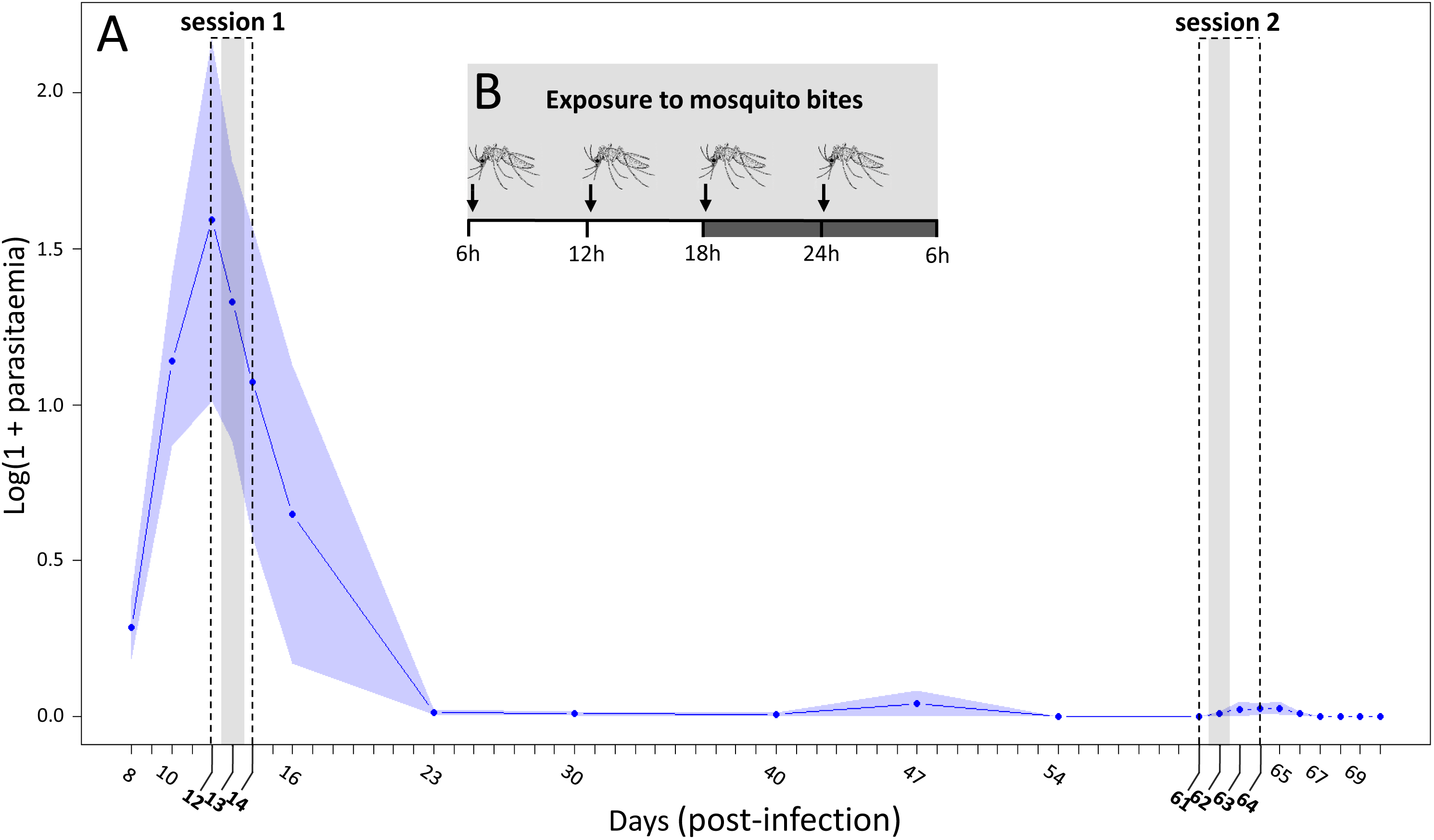
Overview of the experiment with the 2 experimental sessions (grey areas). A) Mean parasitaemia (Log(1 + parasitaemia)), measured at noon, across time post-infection for “unexposed” birds (control). The variation of the parasitaemia among birds is indicated with the shaded envelope (standard error). The dashed boxes represent the two experimental sessions performed in acute (12-14 day post-infection) and in chronic stage (61-64 day post-infection) of infection. In each session, the grey areas correspond to the day at which birds (“exposed”) were exposed to mosquito bites (day 13 post-infection in acute and day 62 post-infection in chronic stage of infection). (B) Zoom on the days where the birds were exposed to mosquito bites. The grey shaded area on x-axis represents the night period. Arrows indicate the time of day at which birds were exposed to mosquito bites. Mosquito exposure were carried out straight after each of the blood sampling events (at 6h, 12h, 18h and 00h).

#### Daily fluctuations of Plasmodium infection

In order to investigate the daily fluctuation of the blood parasitaemia, two experimental sessions were carried out: the first one during the acute stage of infection (Session 1: between day 12 and 14 dpi, **Fig. 1**) and the second one during the chronic stage of infection (Session 2: between day 61 and 64 dpi, **Fig. 1**). During these two experimental sessions blood sampling was carried out every 6 hours (at 6h, 12h, 18h and 00h, **Fig. 1B**). In the acute stage of the infection the existence of a daily fluctuation in the blood parasitaemia was investigated by counting the number of parasites in blood smears (Valkiunas 2004) while in the chronic stage, when parasites in the blood are so scarce that blood smear counts are highly inaccurate, parasite intensities were calculated using molecular tools (see below). In the acute stage of the infection, when the daily fluctuations of parasitaemia may be masked by the large-scale changes in within-host dynamics, the periodicity of the fluctuations in bird parasitaemia was analysed using a new statistical approach that takes into account the overall within-host dynamics of *Plasmodium* infection (see Supplementary Materials, **S1 Text.**).

#### Daily fluctuations of Plasmodium transmission

In order to estimate whether fluctuations in blood parasitaemia translate into fluctuations in transmission to mosquitoes we: 1) obtained estimates of mosquito activity throughout the day, 2) estimated the number of parasites ingested by the mosquitoes at different times during the day and 3) estimated the success of the infection at the oocyst (midgut) stage. For this purpose, on day 13 (Session 1) and day 62 dpi (Session 2), and straight after each of the blood sampling events (at 6h, 12h, 18h and 00h), the birds from the “exposed” treatment were placed inside a cage (L40 x W30 x H30cm) with a batch of 70 uninfected female mosquitoes for 135 minutes. The remaining (“unexposed”) birds were kept under identical conditions but without the mosquitoes. The cages were visited every 45 minutes and all blood fed females were removed and counted. The number of mosquitoes fed at each time step was recorded and was used an as estimate mosquito activity throughout the day (see below). Thereafter, these recently blood-fed mosquitoes were divided in two groups. One half was frozen individually in order to quantify the parasites ingested in the blood meal (see below). The other half was kept alive to obtain an estimate of the blood meal size and of the success of the infection (number of oocysts in the midgut). This was done by placing these mosquitoes in numbered plastic tubes (30 ml) covered with a mesh with a cotton pad soaked in a 10% glucose solution. Seven days later (day 7 post blood meal) the females were taken out of the tubes and the amount of haematin excreted at the bottom of each tube was quantified as an estimate of the blood meal size (Vézilier *et al.* 2010). Females were then dissected and the number of *Plasmodium* oocysts in their midguts counted with the aid of a binocular microscope (Vézilier *et al.* 2010).

At the end of the mosquito exposure session, the parasitaemia of the birds was monitored on a daily basis for a total of 57 days in acute and 8 days in chronic stage of infection. This allowed us to contrast the within-host dynamics of the malaria parasites in birds exposed or not to mosquitoes.

### Molecular analyses

The molecular quantification of parasites in the mosquito blood meal was carried out using a quantitative PCR (qPCR) protocol adapted from (Cornet *et al.* 2013). Briefly, DNA from blood-fed females was extracted using standard protocols (Qiagen DNeasy 96 blood and tissue kit). For each individual, we conducted two qPCRs in the same run: one targeting the nuclear 18s rDNA gene of *Plasmodium* (Primers: 18sPlasm7 5’-AGCCTGAGAAATAGCTACC- ACATCTA-3’, and 18sPlasm8 5’-TGTTATTTCTTGTCACTACCTCTC- TTCTTT-3’), and the other targeting the 18s rDNA gene of the bird (18sAv7 5’ GAAACTCGCAATGGCTCATTAAATC-3’, and 18sAv8 5’-TATTAGCTCTAGAATTACCACAGT TATCCA-3’). All samples were run in triplicate (ABI 7900HT real-time PCR system, Applied Biosystems) and their mean was used to calculate the threshold Ct value (the number of PCR cycles at which fluorescence is first detected, which is inversely correlated with the initial amount of DNA in a sample) using the software Light Cycler 480 (Roche). Parasite intensities were calculated as relative quantification values (RQ). RQ can be interpreted as the fold-amount of target gene (*Plasmodium* 18s rDNA) with respect to the amount of the reference gene (Bird18s rDNA) and are calculated as 2 ^-(Ct18s Plasmodium – Ct18s Bird)^. For convenience, RQ values were standardised by x10^4^ factor and log-transformed (Cornet *et al.* 2013).

### Statistical analysis

The statistical analyses were run using the R software (V. 3.3.3). The different statistical models built to analyse the data are described in the supplementary material (**Table S1**). Analyses where a same individual bird was sampled repeatedly, such as the daily fluctuation of blood parasitaemia or the impact of mosquito exposure on the parasite replication rate, were analysed fitting bird as a random factor into the models (to account for the temporal pseudoreplication), using a mixed model procedure (*lme*, package: nlme). Similarly, mosquito-centred traits (such as infection prevalence or oocyst burden), which may depend on which bird mosquitoes fed on, were also analysed fitting bird as a random factor into the models (to account for the spatial pseudoreplication), using *lme* or *glmer* (package: lme4) according to whether the errors were normally (oocyst burden) or binomially (prevalence) distributed. Time of day and, when necessary, blood meal size (haematin) were used as fixed factors.

Mosquito activity (*i.e.* time required to take a blood meal) was analyzed using survival analyses (package: survival) with time of day (6h, 12h, 18h 00h) fitted as fixed factors in the model and under the assumption of exponential errors. From this model, we estimated the constant hazard rate for each treatment (time of day).

Maximal models, including all higher-order interactions, were simplified by sequentially eliminating non-significant terms and interactions to establish a minimal model (Crawley 2012). The significance of the explanatory variables was established using either a likelihood ratio test (which is approximately distributed as a Chi-square distribution (Bolker 2008) or an F test. The significant Chi-square or F values given in the text are for the minimal model, whereas non-significant values correspond to those obtained before the deletion of the variable from the model. *A posteriori* contrasts were carried out by aggregating factor levels together and by testing the fit of the simplified model using an LRT (Crawley 2012).

To analyse the existence of a circadian rhythm in the parasite dynamics during the acute stage of infection, in addition to the statistical analyses described above, we also developed a new methodology presented in the supplementary materials (**S1 Text**). In addition, we provide a link to a github notebook with a step-by-step description of this procedure and a code that may be used to analyse other within-host time series (https://github.com/QCaudron/timing_malaria_transmission).

## RESULTS

### Theory: evolution of adaptive rhythmicity

To model the evolution of rhythmic transmission strategies we first need to model the epidemiological dynamics of malaria. For the sake of simplicity, the vertebrate host population is assumed to be constant and equal to *N_H_* = *S*(*t*) + *I*(*t*), where *S*(*t*) and *I*(*t*) are the densities of uninfected and infected hosts, respectively. Similarly, the mosquito vector population is also assumed to be constant and equal to *N_v_* = *V*(*t*) + *V*_*I*_(*t*), where *V*(*t*) and *V*_*I*_(*t*) are the densities of uninfected and infected vectors, respectively. The activity of the vector *a(t)* is assumed to fluctuate with a period *T* = 1 day. Low mosquito activity is assumed to decrease biting rate and transmission and, consequently, the epidemiological dynamics also fluctuate periodically. The following set of differential equations describes the temporal dynamics of the different types of hosts (the dot notation indicates differential over time):

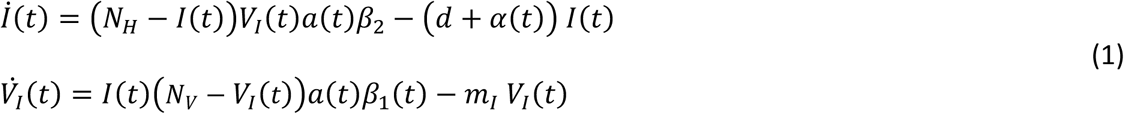

Where *d* is the natural mortality rate of the vertebrate host and *α* is the virulence of malaria (the extra mortality induced by the infection); *m_I_* is the mortality rates of infected vectors; *β*_1_ is the transmission rate from the vertebrate host to the vector; *β*_2_ is the transmission rate from the vector to the vertebrate host.

The pathogen is allowed to have time-varying investment in transmission and virulence in the vertebrate host. As in classical models of virulence evolution, replication allows the parasite to transmit more efficiently (i.e. higher *β*_1_(*t*)) but is assumed to be costly because it may induce the death of the vertebrate host (i.e. higher *α(t))*. To study parasite evolution we track the dynamics of a rare mutant parasite *M* with different transmission and virulence strategies (*β_1*M*_*(*t* and *α_M_*(*t*), respectively):

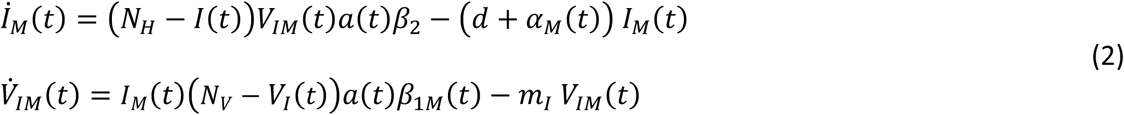

Because the frequency of the fluctuation in mosquito activity is much higher than other dynamical variations of the system we may assume that the density of infected hosts remains approximately stable throughout the day. This separation of time scale allows to focus on the dynamics of the vector compartment which yields:

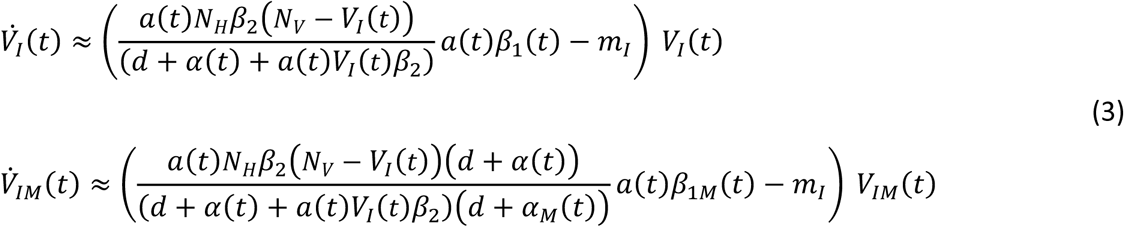

The change in frequency of the mutant is thus given by:

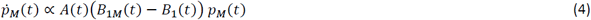

with 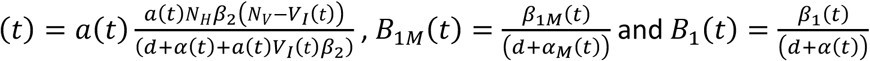

The ability of the mutant to invade the resident population is determined by *s_M_,* the selection coefficient on the mutant, which can be evaluated after integrating the change of the mutant frequency over one day:

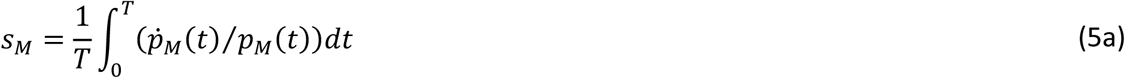

Which yields:

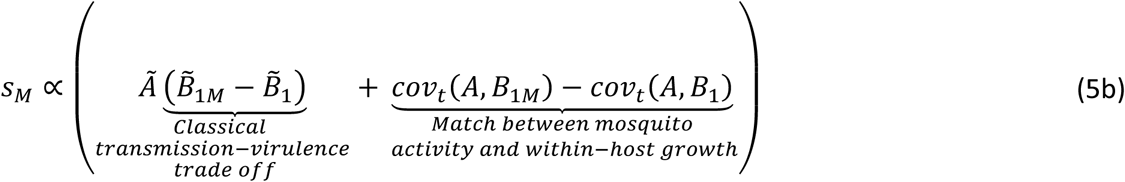

where the tilde refers to the average over a period *T =* 1 day of the fluctuation. The first term in the above equation for *s_M_* is akin to the classical trade-off between transmission *β_1_* and virulence *α.* The second term measures the benefit associated with a closer match between parasite dynamics in the vertebrate host and the rhythmicity in mosquito behavior. For instance, the above expression is particularly useful to examine the invasion of a mutant with a time-varying strategy in a resident pathogen population with a strategy that does not vary with time (i.e. *β_1_* is constant and *cov_t_*(*A,B*_1_) = 0). The mutant will invade only if *s_M_ >* 0 which yields:

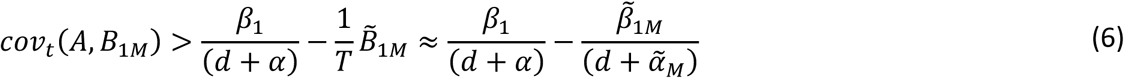

Time-varying transmission thus evolves when the temporal covariance between *A(t),* which is a dynamical variable tightly linked with mosquito activity *a(t),* and investment in transmission is positive and higher than the potential fitness cost (the right-hand side of equation (6)) associated with this time-varying transmission. In other words, this temporal covariance is a measure of the adaptive nature of time-varying transmission.

The above derivation focuses on the evolution of a constitutive time-varying investment in transmission to match fast and periodic fluctuations of vector activity. But when the fluctuation of the environment is slower and/or is less predictable it may be more adaptive to monitor environmental changes and to induce phenotypic modifications accordingly (e.g. Kussell & Leibler 2005). In malaria we developed a similar argument to analyze the evolution of inducible investment in transmission after mosquito bites (Cornet *et al.* 2014).

### Experiment: “Hawking hypothesis” in avian malaria

Blood parasitaemia initially followed a bell-shape function typical of acute *Plasmodium* infections: peaking at day 12 post-infection (dpi) and decreasing thereafter (**Fig. 1**). The infection subsequently entered a long-lasting chronic state, which was characterized by a low blood parasitaemia over several weeks (**Fig. 1**). During the acute phase of the infection, and before the exposure to the mosquitoes, there was no significant difference in the parasitaemia of the hosts assigned to the “exposed” and “unexposed” treatments (model 1: *χ*^2^1= 0.01, p = 0.941, **Fig. 2A**). However, after they had been exposed to the mosquito bites, the acute-phase parasitaemia of the “exposed” birds was significantly higher than that of their “unexposed” counterparts (model 2: *χ*^2^1= 8.59, p = 0.003, **Fig. 2A**). This effect was short-lived and only lasted around 48h (peak reached in 24h, **Fig. 2A**). During the chronic phase of the infection, there was a significant difference in the parasitaemia of the birds before the exposure session (model 3: *χ*^2^1= 10.83, p = 0.001, **Fig. 2B)**: “unexposed” birds had a higher parasitaemia than “exposed” hosts. After exposure to mosquito bites, while the parasitaemia of the “unexposed” birds did not vary (model 4: *χ*^2^1 = 0.086 p = 0.771, **Fig. 2B**), the parasitaemia of the “exposed” chronically-infected hosts increased over time (peak reached in 6 days, model 5: *χ*^2^1 = 22.99 p < 0.0001, **Fig. 2B**).

**Figure 2:**
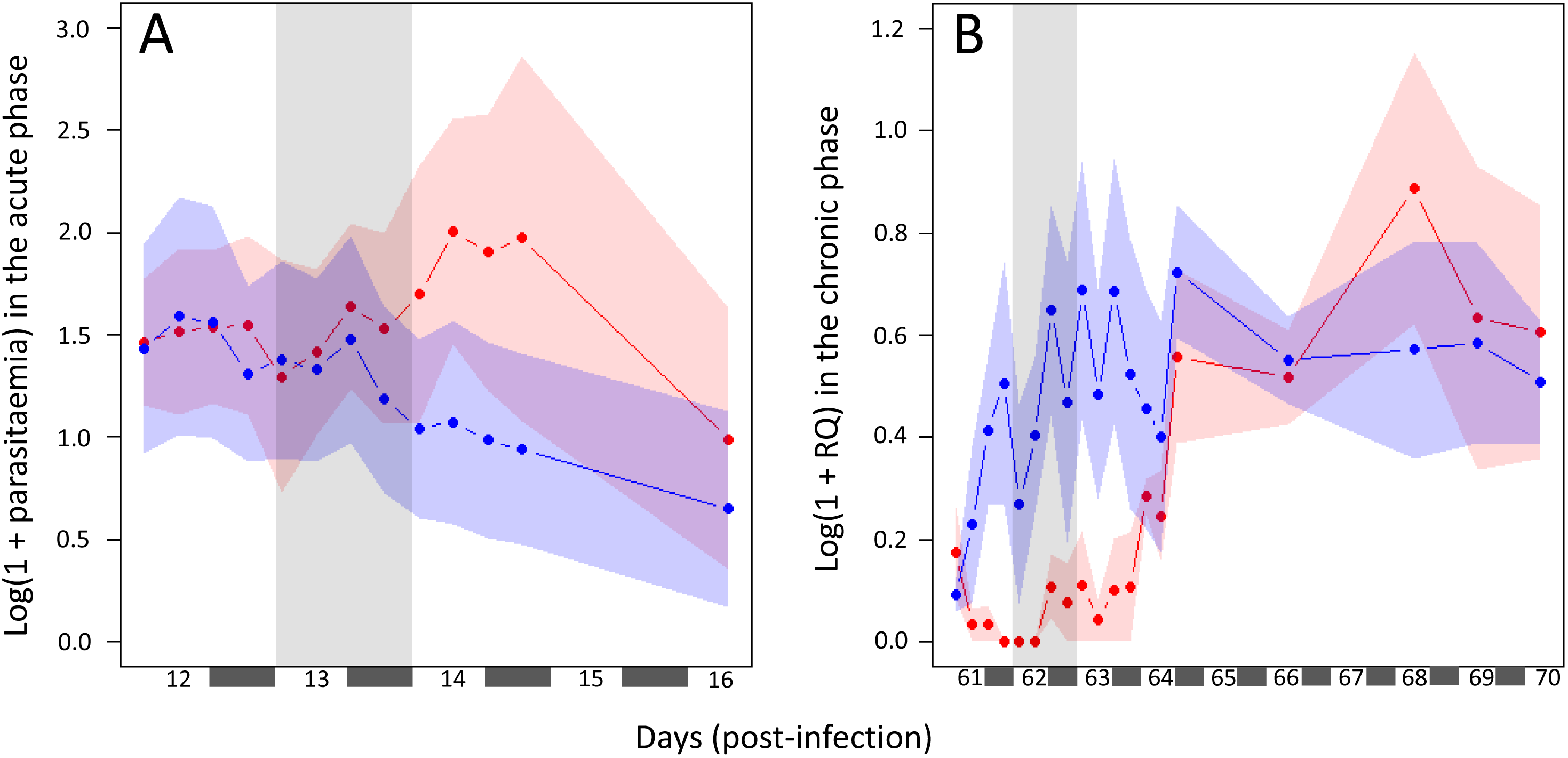
Within-host dynamics of blood parasitaemia (mean ± se) of *Plasmodium relictum* in birds. The dynamics in birds exposed or unexposed to mosquito bites is represented in red and blue, respectively. Mosquito exposure took place (A) day 13 (at 6AM, 12AM, 6PM and 12PM) and (B) day 62 (at 6AM, 12AM, 6PM and 12PM) post-infection.

### Daily fluctuations of blood parasitaemia

The periodicity of the fluctuations in bird parasitaemia was explored using a new statistical approach that takes into account the overall within-host dynamics of *Plasmodium* infection during the acute phase of the infection (See **Supporting Information**). In spite of a limited number of samples this analysis suggests that bird parasitaemia fluctuates periodically with a peak in the late afternoon (See **Supporting Information**). We then examined in depth these daily fluctuations in parasitaemia and their consequences on mosquito transmission in both the acute and chronic phases of the infection. To avoid the potential effect of mosquito bites on within-host dynamics, we focused our analyses on “exposed” birds. In the acute phase of the infection we found a significant effect of the time of day on blood parasitaemia (model 6: *χ*^2^1 = 11.58, p = 0.009, **Fig. 3A**). The parasitaemia was highest in the evening (18h) and lowest early in the morning (6h, **Fig. 3A**). During the chronic phase of the infection blood parasitaemia was very low in all exposed birds (parasitaemia < 0.001%, **Fig. 1**). Molecular methods, however, allowed us to detect daily variations in parasitaemia. Parasite burden was null at 6h and 12h, or below the detection levels, but increased in the evening (**Fig. 3B**).

**Figure 3:**
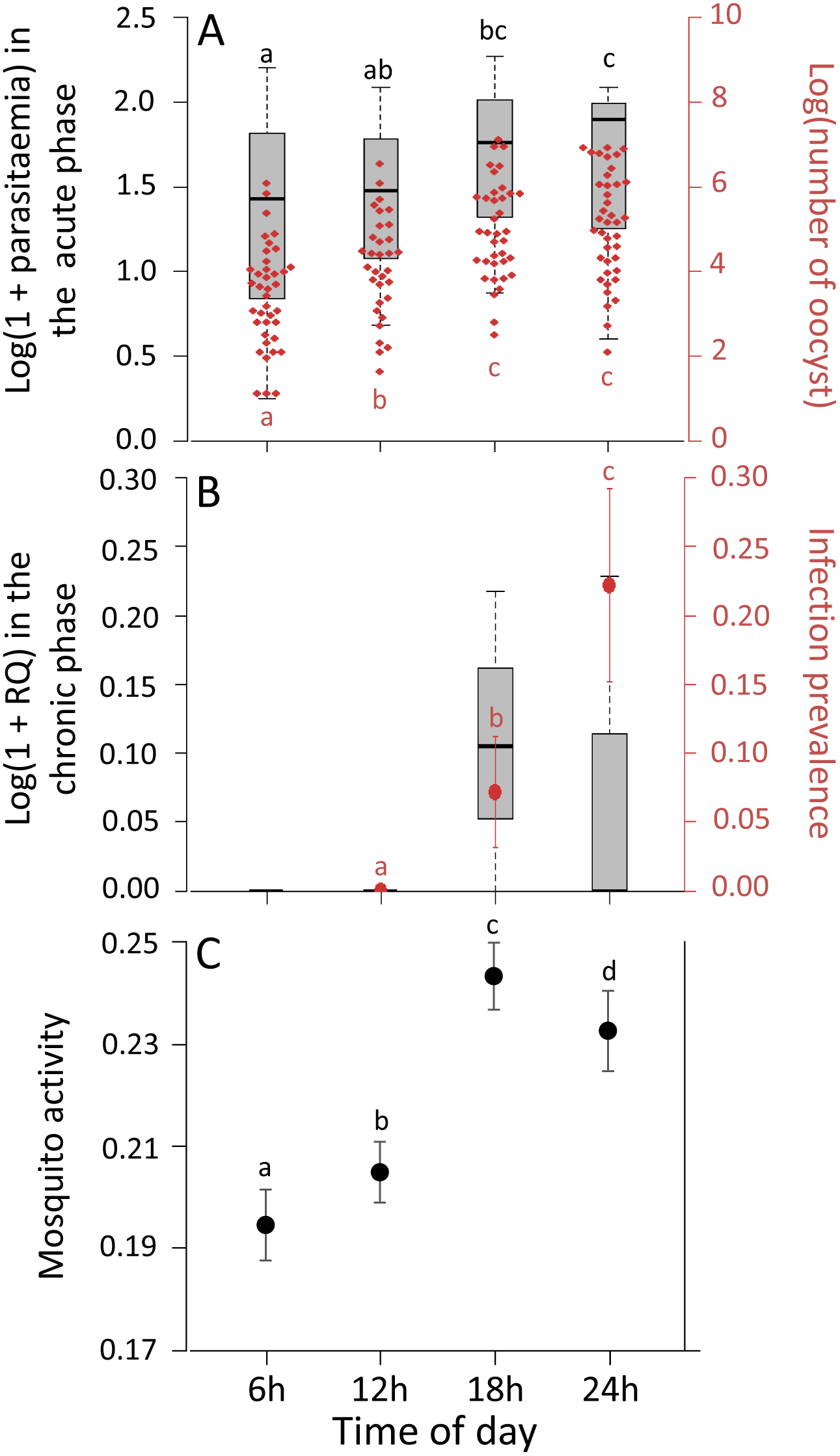
Timing of malaria within-host dynamics and mosquito activity in avian malaria. A) Daily fluctuations of *Plasmodium* transmission in acute phase of infection (session 1: day 13 post-infection, see **Fig. 1**). Boxplot represent the blood parasitaemia (Log(1 + parasitaemia)) of the exposed birds measured at 6h, 12h, 18h and 00h, 13 days after the infection by *Plasmodium*. The red points represent the distribution of the number of oocysts in the midgut of *Plasmodium*-infected females 7 days after the blood meal. Blood meals were taken on the birds whose parasitaemia is described by the boxplots. (B) Daily fluctuations of *Plasmodium* transmission in chronic phase of infection (session 2: day 62 post-infection, see **Fig. 1**). Boxplot represent the blood parasitaemia (Log(1 + Relative Quantification values)) of the exposed birds measured at 6h, 12h, 18h and 00h, 62 days after the infection by *Plasmodium*. The red points represent the prevalence of *Plasmodium* infection in females 7 days after the blood meal. Blood meals were taken on the birds whose parasitaemia is described by the boxplots. (C)Daily fluctuations of mosquitoactivity.From the survival analyses (see Materials and Methods), the constant hazard rate for each treatment (time of day) and the standard errors were calculated. Levels not connected by same letter are significantly different.

### Daily fluctuations of *Plasmodium* transmission

As mentioned above, the aim of this section was to investigate whether fluctuations in blood parasitaemia translate into fluctuations in transmission to mosquitoes. For this purpose, we first quantify the number of oocysts in mosquitoes fed at different times of the day. We then explore whether these differences can be explained by differences in the amount of parasites ingested by the mosquitoes at different times of the day. Finally, we explore whether the fluctuations in blood parasitaemia and mosquito infectivity match the daily patterns of mosquito activity.

In the acute stage, the mosquito infection prevalence was 100% for all feeding times. Blood feeding time, however, had a very significant effect on the oocyst burden of mosquitoes (model 7: *χ*^2^1 = 42.69, p < 0.0001, **Fig. 3A**). Females that fed in the evening (18h and 00h) had more than twice as many oocysts as those feeding at noon (contrast analyses: 12h/18h:*χ*^2^1 = 8.28, p = 0.004, 12h/00h:*χ*^2^1 = 13.92, p < 0.0001, oocysts burden: 12h: mean ± s.e: 108 ± 25, 18h: 262 ± 51 and 00h: 314 ± 52) and noon-feeding mosquitoes had significantly more oocysts than those feeding in the early morning (contrast analyses: 6h/12h:*χ*^2^1 = 5.03, p = 0.025, oocysts burden: 6h: 62 ± 14). As expected, haematin, a proxy for blood meal size, has an impact on mosquito oocyst burden (model 7: *χ*^2^1 = 49.17, p < 0.0001). Crucially, however, the time of day has no impact on haematin production (model 8: *χ*^2^1 = 6.54, p = 0.091) implying that the blood meal sizes do not change according to the feeding times. An impact of bird parasitaemia on oocyst burden was observed but only when the feeding time was removed from our statistical model (model 7: with time of day as covariate *χ*^2^1 = 1.70, p = 0.192, model 9: without time of day as covariate *χ*^2^1 = 15.09, p < 0.001).

The quantification of parasites ingested by mosquitoes showed a significant positive correlation with both haematin and time of day (model 10: *χ*^2^1 = 28.01, p < 0.0001, χ²1 = 41.71, p < 0.0001 respectively). The quantity of parasite ingested by mosquito was highest at midnight (00h) and lowest early in the morning (6h). Bird parasitaemia also had an impact on the quantity of parasites ingested by females but only when the time of day was removed from the statistical model (model 10: with time of day as covariate *χ*^2^1 = 0.12, p = 0.727, model 11: without time of day as covariate *χ*^2^1 = 3.62, p = 0.047).

In the chronic stage of the infection, mosquito infection prevalence varied throughout the day (model 12: *χ*^2^1 = 6.98, p = 0.030, **Fig. 3B**). Infection prevalence was 0% at noon, 7% (mean ± s.e: 7.1 ± 4) at 18h and 22% (22.2 ± 7.1) at 00h (no data available for 6h, contrast analyses: 12h/18h:*χ*^2^1 = 3.89, p = 0.049, 12h/00h:*χ*^2^1 = 6.34, p = 0.012, 18h/00h:*χ*^2^1 = 3.91, p = 0.048). However, oocyst numbers were too low (all infected females had a single oocyst) to detect any effect of time of day on parasite burden. Bird parasitaemia and blood meal size (haematin) had no impact on mosquito infection prevalence (model 12: with time of day as covariate *χ*^2^1 = 0.58, p = 0.447, model 13: without time of day as covariate *χ*^2^1 = 1.64, p = 0.201, model 12: with time of day as covariate *χ*^2^1 = 0.64, p = 0.725, model 13: without time of day as covariate *χ*^2^1 = 0.17, p = 0.679 respectively).

### Daily fluctuations of mosquito activity

Mosquito activity was significantly impacted by the time of day (model 14: *χ*^2^1 = 204.15, p < 0.0001, **Fig. 3C**). Overall, the activity of vectors was higher in the evening (18h, 00h) than in the morning (6h, 12h). The maximal activity was observed at dusk (contrast analyses: 18h/00h: *χ*^2^1 = 28.78, p < 0.0001, 18h/12H: *χ*^2^1 = 148.89, p < 0.0001, 18h/6H: *χ*^2^1 = 166.48, p < 0.0001, **Fig. 3C**) and the minimal activity at dawn (contrast analyses: 6h/12h: *χ*^2^1 = 4.09, p = 0.026, 6h/00h: *χ*^2^1 = 38.90, p < 0.0001, **Fig. 3C**). Interestingly, these daily variations in mosquito activity were positively correlated with both bird parasitaemia and parasite transmission to mosquito in acute (**Fig. 4A**) but also in chronic stage of infection (**Fig. 4B**).

**Figure 4:**
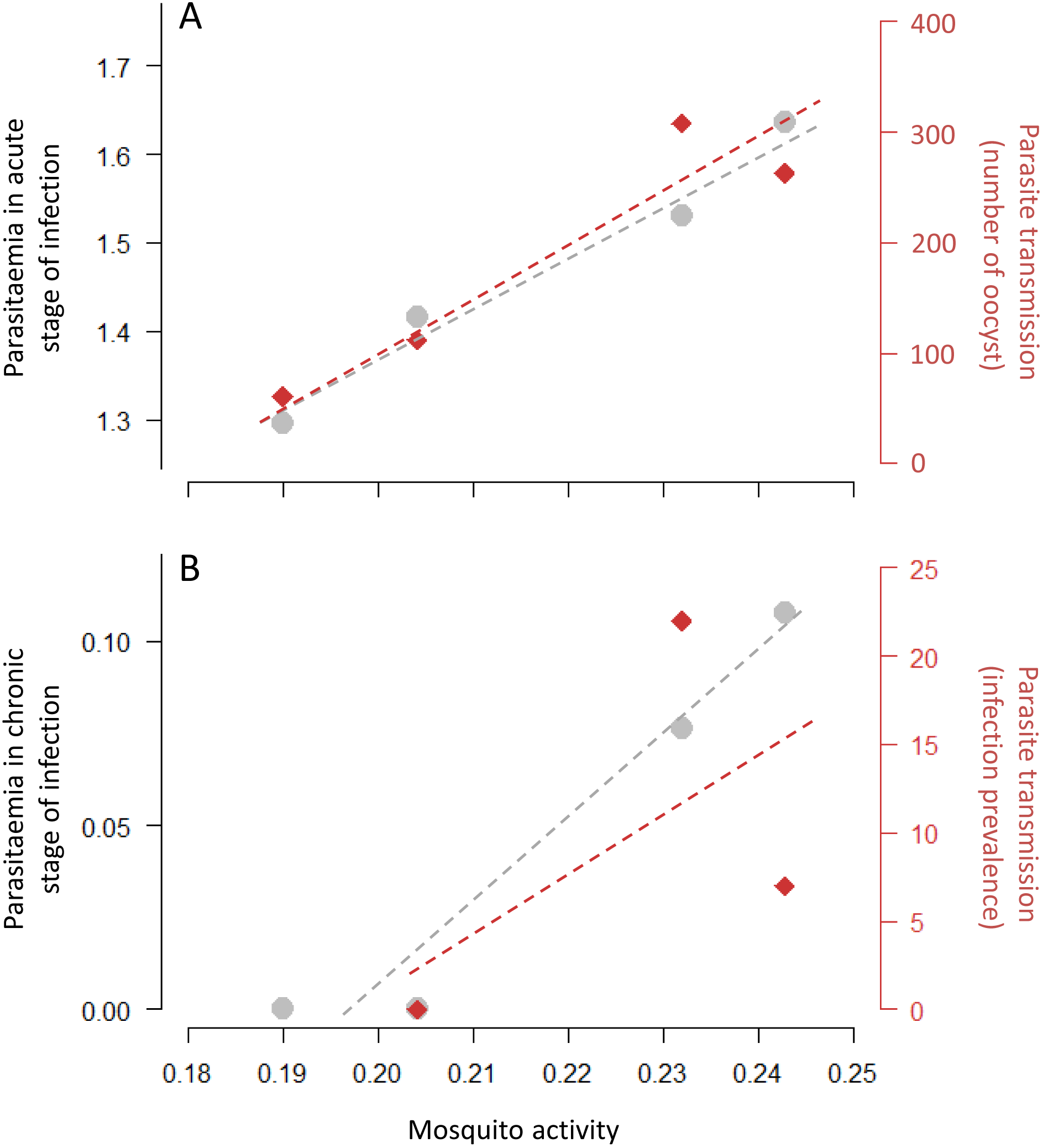
Testing the “Hawking hypothesis” in avian malaria. A) Correlation between mosquito activity (constant hazard rate estimated from model 14, **TableS1**) and bird parasitaemia (log(1+parasitaemia), in grey) and parasite transmission to mosquito (infection intensity: oocyst burden, in red) in acute stage of infection. (B) Correlation between mosquito activity (constant hazard rate estimated from model 14, **TableS1**) and bird parasitaemia (log(1 + Relative Quantification values), in grey) and parasite transmission to mosquito (infection prevalence (%), in red) in chronic stage of infection.

## DISCUSSION

Temporal fluctuations of the activity of mosquito vectors have profound consequences on malaria transmission (Barrozo *et al.* 2004; Lalubin *et al.* 2013). Here we argue that *Plasmodium* parasites have evolved two different and complementary transmission strategies to cope with these variations of their environment: a constitutive time-varying strategy that generates a covariance between parasite investment in transmission and vector activity and a plastic, fast acting, strategy that allows the parasite to react rapidly to the presence of mosquitoes.

First, our theoretical model indicates that fast and predictable oscillations in mosquito activity can select for a constitutive time-varying strategy in the parasite, provided this strategy generates a positive covariance between the activity of the vector and the parasite’s investment in transmission (see equation (6)). Our experimental results show both that the activity of *Culex* mosquitoes oscillates throughout the day in a predictable way (**Fig. 3C**) but also, that these daily fluctuations of mosquito activity are matched with periodic fluctuations in malaria transmission during both phases of the infection *(i.e.* acute and chronic, **Fig. 4**). This positive covariance supports the “Hawking hypothesis” and the idea that this time-varying transmission may result from an adaptation of the pathogen.

Second, our experiment demonstrates the existence of plastic transmission strategies enabling avian malaria parasites to respond to mosquito bites. In a previous study, we showed that mosquito bites stimulate within-host growth and investment in transmission during the chronic phase of *Plasmodium relictum* infections (Cornet *et al.* 2014). In the present study, we obtain a similar effect in the chronic but also in the acute phase of the infection. This plastic transmission strategy is expected to evolve when variations in the abundance of their mosquito vectors are less predictable (Cornet *et al.* 2014; Reece & Mideo 2014). During the chronic phase of the infection, such plastic transmission strategies may allow the parasite to react to the seasonal variations in mosquito abundance and to reactivate its transmission when mosquitoes are around (Cornet *et al.* 2014; Reece & Mideo 2014). During the acute phase of the infection, this strategy may also allow the parasite to respond to unexpected variations in the abundance of mosquitoes driven by stochastic processes such as variations in temperature and humidity (Yamana & Eltahir 2013).

In spite of the match between these theoretical predictions and our experimental results, our adaptive hypothesis is challenged by alternative explanations for the existence of periodic variations in parasitaemia and mosquito infection. Several studies suggest that the dynamics of the infectivity of *Plasmodium* might not be underpinned by the feeding activity cycle of its vector but induced by the vertebrate immunity (see Mideo *et al.* 2013), whose activity is known to vary during the day (Scheiermann *et al.* 2013; Curtis *et al.* 2014). This variation may alter the number and/or the infectiousness of gametocytes and explain (at least partly) the increase of transmissibility during the evening. It would be interesting to monitor whether the efficacy of the birds’ immune system to fight against a *Plasmodium* infection fluctuates throughout the day, and to evaluate its potential effect on the transmissibility of avian malaria.

In addition, the increase in mosquito infection may also be explained by physiological cycles in the vector. Daily cycles in the production of immune compounds (Rund *et al.* 2016; Tsoumtsa *et al.* 2016) or molecules (*e.g.* nutrients) used by *Plasmodium* (Carter *et al.* 2007; Dinglasan *et al.* 2007) may impact the viability of ookinetes or their ability to invade the midgut epithelia. One way to quantify this effect would be to perform similar experiments with vectors with the circadian rhythm experimentally inversed (jet-lagged). Reversed patterns of time-varying infectivity in jet-lagged and control mosquitoes would indicate a strong effect of the circadian rhythm of the insect vector. In contrast, if both jet-lagged and control mosquitoes exhibit similar patterns of infection this would indicate that the infectivity is under the parasite’s control and would support the “Hawking hypothesis”.

The most efficient way to demonstrate unequivocally the adaptive nature of these time-varying transmission strategies would be to perform experimental evolution (Johnson 2005). For instance, does the parasite lose its ability to react to mosquito bites if the parasite is always transmitted from bird to bird by intraperitoneal injection (Pigeault *et al.* 2015)? Could the parasite be made to evolve other patterns of daily investment in transmission if the mosquitoes are allowed to feed on birds at very specific time of the day? Avian malaria provides a perfect experimental system to carry out such experiments. Earlier studies have observed a great degree of variation in the period and in the phase of the fluctuations of within-bird dynamics. For instance, *P. circumflexum* has a periodicity of 48h peaking in the late afternoon, while *P. elongatum*’s periodicity is 24h and peaks in the early morning (see Hewitt 1940 for a review). Besides, the amplitude of the fluctuations of parasitaemia reported in some of these earlier experimental studies is orders of magnitude higher than the one we observed in the present study (Taliaferro 1925, Huff & Bloom 1935, Hewitt 1940). What factors explain the maintenance of such a large amount of natural variation? Additional experimental studies using different avian *Plasmodium* lineages would yield unique perspectives on the adaptive nature of the rhythmicity of malaria within-host dynamics. The genomic analysis of evolved lines would also yield new candidate genes governing these key adaptations. This deeper understanding of malaria transmission may thus yield practical implications for the control of human malaria parasites.

## Data archiving

Data for this study will be made available once the manuscript accepted for publication (Dryad website)

## Authors’ contributions

Conceived and designed the experiments: RP AR SG. Performed the experiments: RP AN. Analysed the data: RP. Developed and analysed the theoretical model: SG. Developed the new statistical methodology to study daily fluctuations of parasitaemia: QC. Wrote the paper: RP AR SG. All authors read and commented the paper and approved the final version of the manuscript.

